# Differences in gait stability and acceleration characteristics between healthy young and older females

**DOI:** 10.1101/2021.02.24.432667

**Authors:** Yuge Zhang, Xinglong Zhou, Mirjam Pijnappels, Sjoerd M. Bruijn

**Author notes:** **Corresponding:** Sjoerd M. Bruijn.

## Abstract

Our aim was to evaluate differences in gait acceleration intensity, variability and stability of feet and trunk between older females and young females using inertial sensors. Twenty older females (OF; mean age 68.4, SD 4.1 years) and eighteen young females (YF; mean age 22.3, SD 1.7 years) were asked to walk straight for 100 meters at their preferred speed, while wearing inertial sensors on heels and lower back. We calculated spatiotemporal measures, foot and trunk acceleration characteristics and their variability, as well as trunk stability using the local divergence exponent (LDE). Two-way analysis of variance (including the factors foot and age), Student’s t-test, and Mann–Whitney U test were used to compare statistical differences of measures between groups. Cohen’s d effects were calculated for each variable. Foot maximum vertical acceleration and amplitude, trunk-foot vertical acceleration attenuation, as well as their variability were significantly smaller in OF than in YF. In contrast, trunk mediolateral acceleration amplitude, maximum vertical acceleration, and amplitude, as well as their variability were significantly larger in OF than in YF. Moreover, OF showed lower stability (i.e., higher LDE values) in mediolateral acceleration, mediolateral and vertical angular velocity of the trunk. Even though we measured healthy older females, these participants showed lower vertical foot accelerations with higher vertical trunk acceleration, lower trunk-foot vertical acceleration attenuation, less gait stability, and more variability of the trunk, and hence, were more likely to fall. These findings suggest that instrumented gait measurements may help for early detection of changes or impairments in gait performance, even before this can be observed by clinical eye or gait speed.

## 1 Introduction

Falls among older adults are the leading indirect cause of disability and death (Tang et al., 2017). Epidemiological studies have shown that the 30% of people aged 65 years and older fall, with an increase in incidence to 40% in people over 80 years (Weber et al., 2018). This is due to poorer physiological function and control of stability with ageing (Winter, 1995). In China, 53% of falls occur while walking (Xia et al., 2010), and hence, it is particularly important to pay attention to gait performance of older adults for early identification of stability problems to prevent falls. Moreover, many studies have shown that among people over 60 years, females were more likely to fall (Tinetti et al., 1988;Daley and Spinks, 2000;De Rekeneire et al., 2003), as about 65% of women and 44% of men fell in their usual place of residence (Masud and Morris, 2001). Therefore, we focused on gait stability of females in our study.

There are several ways to evaluate gait, such as clinical function tests, questionnaires and measurements in a biomechanics laboratory (Hamacher et al., 2011). Questionnaires and clinical tests cannot reflect gait performance outside the laboratory, and sometimes have poor objectivity (van Schooten et al., 2015a). Gait assessment in a biomechanical laboratory has the advantage of capturing whole body kinematics which is accurate but also costly, time-consuming and limited to space and time (Terrier and Deriaz, 2011). Nowadays, the feasibility of inertial sensor to quantify the whole-body gait kinematics has been demonstrated(Tao et al., 2012), and it can be used to collect gait data in people’s own environment by a single sensor on either the trunk or foot (Zhu et al., 2012;Weiss et al., 2013).

Gait stability reflects the ability to keep walking in the face of perturbations (Pai and Bhatt, 2007;Bruijn et al., 2013). Dynamical systems and non-linear time series analysis can be used to evaluate gait stability by quantifying the complex and chaotic characteristics of the human body (Bressel, 2004). One of these measures, the local divergence exponent (LDE) has been shown to have good reliability and validity (England and Granata, 2007;Son et al., 2009;Hu et al., 2012). The LDE quantifies the average exponential rate of divergence of neighboring trajectories in state space, and provides a direct measure of the sensitivity of a system to small perturbations (Dingwell and Marin, 2006).

Internal perturbations of the human body cause variability and randomness in gait (Zhang et al., 2011). If gait is within a stable range, people would not need to correct this variability. Increased variability likely reflects a less automatic gait pattern, instability and increased susceptibility to falls (Weiss et al., 2013). Studies also confirmed that variability in some gait characteristics (such as stride length, stride width, stride time) is highly related to the risk of falling (O’Loughlin et al., 1994;Chau et al., 2005). However, some studies suggested that variability is not equal to stability, as the level of variability was not necessarily negatively related to the level of stability (Li et al., 2005;van Emmerik et al., 2016).

As the control of stability in gait declines with ageing, we aimed to use inertial sensors to assess differences in gait stability and variability between healthy young and older females. In doing so, we focused on data obtained from trunk as well as foot sensors and calculated acceleration intensity, stability, and variability measures. We hypothesized that older females have a lower gait stability and increase variability on trunk accelerations compared with younger females.

## 2 Materials and Method

### 2.1 Participants

A total of 20 healthy older females (OF) and 18 younger females (YF) were recruited from the campus of Beijing Sport university, China (Table 1). None of our participants had any orthopedic or neurological disorders, acute pain or other complaints that might have affected gait and they were all able to walk independently without a walking aid. All participants were informed about the research procedures and the protocol was approved by the Ethics Committee of Sports Science Experiment of Beijing Sport University (approval number: 2021010H).

**Table 1.** Participant characteristics

| Groups | YF | OF | p-Value (T-test) |
| --- | --- | --- | --- |
| Age (years) | 22.3 (1.7) | 68.4 (4.1) | <0.001*** |
| Height (m) | 1.65 (0.04) | 1.59 (0.05) | <0.001*** |
| Body mass (kg) | 54.66 (3.93) | 63.2 (7.95) | <0.001*** |
| BMI (kg/m<sup>2</sup>) | 20.19 (1.53) | 24.96 (2.60) | <0.001*** |
| Leg length (cm) | 88.21 (3.53) | 87.28 (3.19) | 0.29 |

### 2.2 Data Acquisition

Participants wore three inertial sensors (Xsens MTw Awinda, the Netherland) on heels and on the lumbar region of the trunk, using the supplied elastic belt. These sensors had a sample rate of 100 samples/s and a range of -160 m/s^2^ and +160 m/s^2^. Data collection was synchronized between sensors. All participants wore the same model of shoes. They were asked to walk 100 meters on a straight running track at a self-selected speed, since gait variability is expected minimal at this speed for healthy people (Hausdorff et al., 1997). In addition, although clinical gait tests are usually 4 or 10 meters, these tests do not represent daily-life gait very well (Van Ancum et al., 2019). Therefore, 100 meters used in this study can reflect well the natural gait at a comfortable speed without participants being exhausted.

### 2.3 Gait Measures

MATLAB (R2019b, MathWorks Inc, Natick, MA, USA) was used to analyze data without the first and last steps. Each gait cycle was identified from the sagittal plane angular velocity of foot sensors with three gait events: heel-strike (T_heel_strike_), toe-off (T_toe_off_) and foot-flat (T_foot_flat_) (Mariani et al., 2010). Stride time was defined as the duration between two consecutive T_heel_strike_. Combined with the gait events of both feet, we got the initial double support period (IDS) and the terminal double support period (TDS).

For the trunk sensor, sensor data were realigned to a coordinate system based on the accelerometer’s orientation with respect to gravity (vertical axis) and optimization of left-right symmetry (mediolateral axis) (Rispens et al., 2015;Van Schooten et al., 2015b).

For the foot sensors, initial displacements were calculated by integrating linear accelerations twicer for each gait cycle (in the global coordinate system), using zero-velocity-update method to eliminate drift, assuming linearity of the drift (Skog et al., 2010). The hence obtained direction of displacement was not necessary along the x or y axis of the global coordinate system. To obtain meaningful stride lengths, we thus rotated the obtained positions, the acceleration and angular velocity of the feet to a coordinate system that was aligned with the direction of walking (i.e., end position minus starting position), with the vertical axis being vertical. Then, walking speed was obtained by dividing the distance of the walking direction by the time.

For acceleration measures, maximum vertical acceleration of feet and trunk were calculated to reflect the intensity of ground contact (Gill and O’Connor, 2003). It has been suggested that people stabilize their head during walking (Kavanagh et al., 2006). Although the trunk segment plays a key role in damping gait-related oscillations (Kavanagh et al., 2006), the damping of oscillations by the trunk in the vertical direction has been suggested to be minor (Prince et al., 1994;Kavanagh et al., 2004). Hence, such accelerations must be attenuated by the lower limbs. Thus, we calculated trunk-foot vertical acceleration attenuation was used in our study, which was calculated by the difference in maximum vertical acceleration between trunk and foot, which represents the impact absorption of the lower limbs. Acceleration amplitude (in the coordinate system prescribed by the walking direction, see above) for each direction (anteroposterior direction (AP), mediolateral (ML) and vertical (VT)) was calculated as the range of acceleration in a gait cycle.

For above measures of each person, after getting the mean and standard deviation (SD) over all cycles (see Table 2, Table3), we obtained coefficient of variation (CV) by dividing the standard deviation by the mean (Kavanagh and Menz, 2008) (see Supplementary Table 1).

**Table 2.**
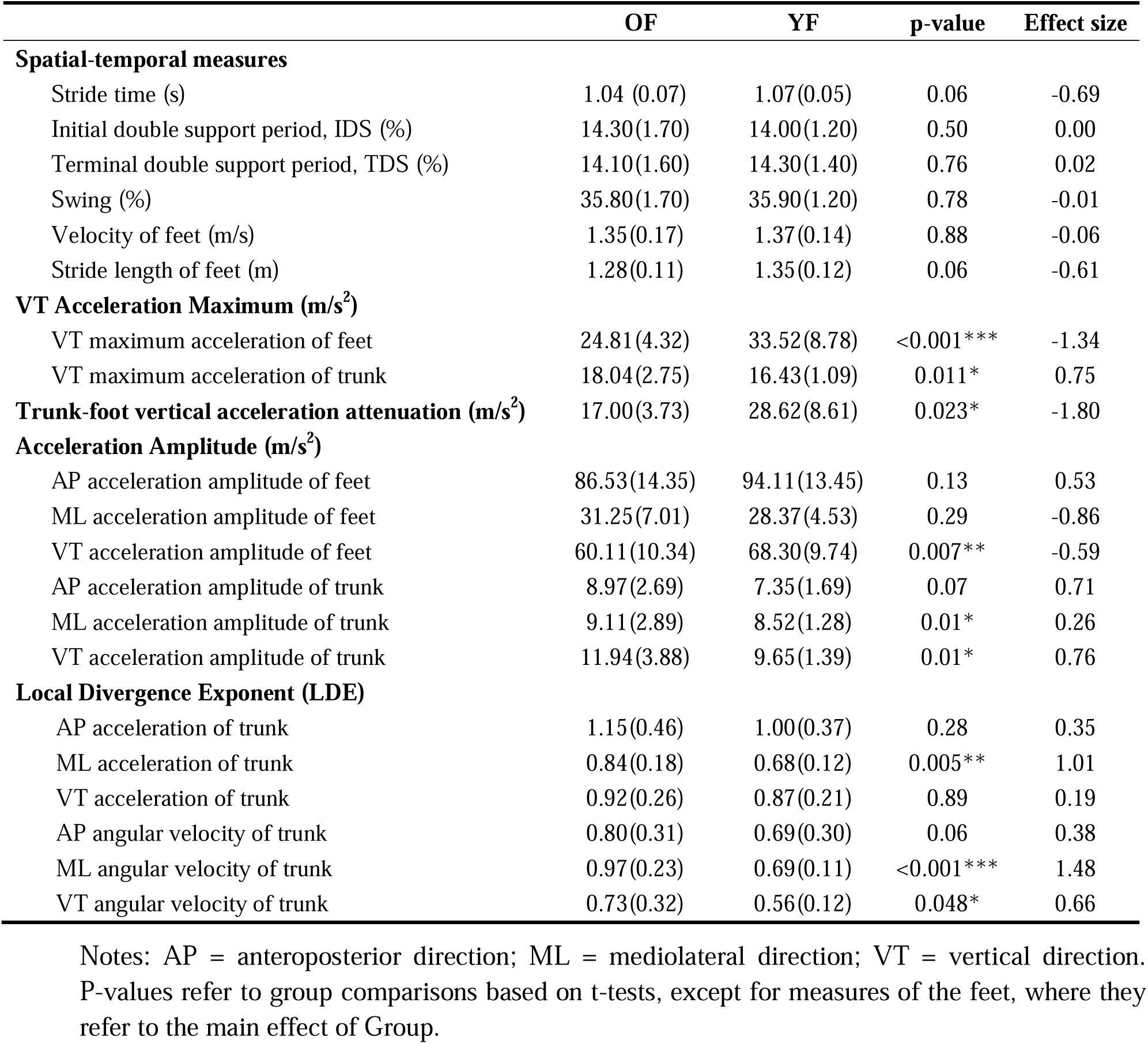
Mean (and SD) of all gait measures

**Table 3.**
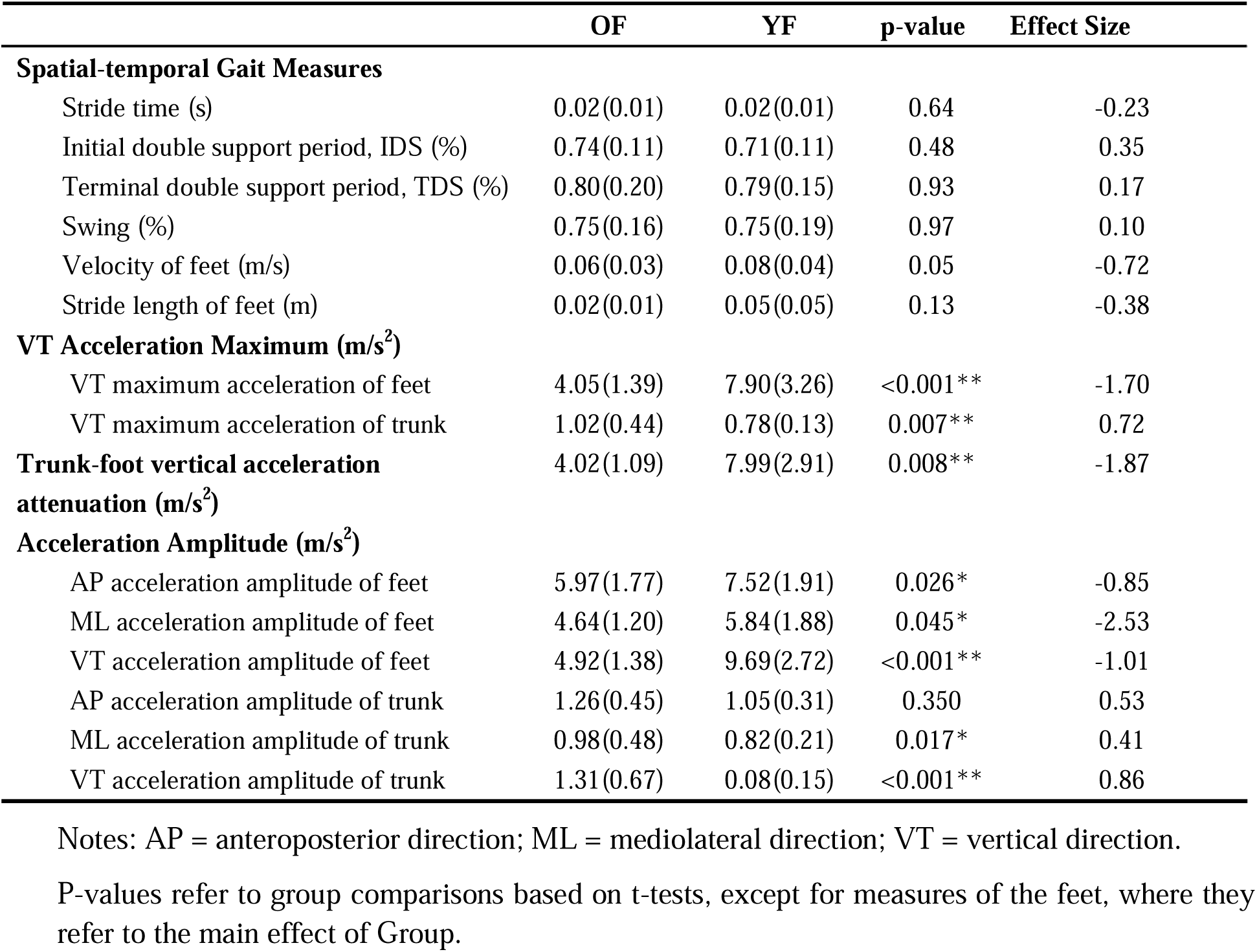
Variability (and SD) of all gait measures

We calculated the LDE of acceleration and angular velocity of each dimension separately (in the coordinate system prescribed by the walking direction, see above). The time series of 50 gait cycles was normalized into 5000 samples, with the average of 100 samples per cycle. From these data, state spaces were reconstructed using the method of correlation integral (C-C method), which not only can determine both embedding dimension and delay time, but also has a good robustness to the noise in small amount of data (Kim et al., 1999) (see Supplementary Tables 2 and 3 for dimension and delay values). LDE was expressed as the mean logarithmic rate of divergence per stride using Rosenstein’s method (Rosenstein et al., 1993). Higher values of the LDE indicate a lower local stability.

### 2.4 Statistical Analysis

Normality was assessed using the Kolmogorov–Smirnov test. For measures of the left and right feet, differences were tested using two-way ANOVAs, with within-subject factor Foot (left and right) and between-subject factor Group (YF and OF). For other measures, we used Student’s t-tests to compare between age groups. For LDE, which appeared not distributed normally, we compared between groups using the Mann–Whitney U test. For all measures, p<0.05 was considered as a significant effect. Cohen’s d effects were calculated for each variable as the difference between group means divided by the group pooled standard deviation. Magnitudes of d = 0.01, 0.20, 0,50, 0.80, 1.20 and 2.0 were considered very small, small, medium, large, very large and huge, separately (Sawilowsky, 2009;Charach et al., 2011;Cohen, 2013).

## 3 Results

Descriptive characteristics of the participants are summarized in Table 1. OF were significantly older, shorter and had a higher weight and higher BMI than YF. The mean age of OF and YF was 68.4 and 22.3, respectively.

Table 2 shows the mean values for all measures. We found no interaction between Foot and Group for any of the outcome measures, and no significant effect of Foot. Hence, all variables that were calculated for both feet are displayed as averages over both feet. OF had higher maximum vertical acceleration of the trunk than YF, with medium effect size (0.75), but smaller maximum vertical acceleration of the feet than YF, with very large effect size (1.34). As a result, OF had significantly smaller trunk-foot vertical acceleration attenuation, with a very large effect size of 1.8. In addition, OF’s vertical accelerations amplitude of the feet were significantly smaller than YF, with medium effect size (−0.59). For the trunk, OF’s ML and VT acceleration amplitude were significantly larger than YF, and the effect size of the latter was largest (0.76). The LDE of trunk from ML acceleration, and from ML and VT angular velocity were significantly larger (less stable) for OF than for YF, with large (1.01), very large (1.48) and medium effect size (0.66), respectively.

Table 3 shows the variability of all measures. No significant difference in variability of spatial-temporal gait measures were found between groups. The variability of maximum vertical acceleration of the feet was significantly smaller for OF than YF, and its effect size was 1.70. While for trunk, the variability of the maximum vertical acceleration was significantly larger for the OF (medium effect size 0.72). The variability of trunk-foot vertical acceleration attenuation was smaller in OF than in YF, (effect size very large, 1.87). OF had significantly smaller variability of acceleration amplitude of the feet in three directions than YF, with huge effect size in ML direction (2.53) and a large effect size in the VT direction (1.01). For the trunk, OF’s variability of acceleration amplitude was significantly larger than YF in ML and VT direction, with effect sizes of 0.41 and 0.86, respectively. The CV of gait measures showed largely the same pattern as the SD (see Supplementary Table 1).

## 4 Discussion

### 4.1 Mean gait measures

In this study, we used inertial sensors to evaluate differences in acceleration intensity, variability and stability of feet and trunk during gait between healthy young and older females. Although older adults generally were suggested to walk slower due to physical limitations like muscle weakness or loss of flexibility (Hamacher et al., 2014), the OF in our study walked at similar preferred speed and stride length as the YF.

We found a reduction in foot vertical maximum acceleration in OF, which probably reflected a reduction of peak ground reaction forces. Such a reduction of ground reaction forces could result from a crouch-like gait, which has been shown in young adults to lead to a reduction of the peak ground reaction force (Li et al., 1996;Grasso et al., 2000). Such a crouch like gait may increase the metabolic cost of locomotion in the elderly (Carey and Crompton, 2005). Although the trunk segment plays a key role in damping gait-related oscillations (Kavanagh et al., 2006), the damping of oscillations by the trunk in the vertical direction has been suggested to be minor (Prince et al., 1994;Kavanagh et al., 2004). In our study, we found a lower trunk-foot vertical acceleration attenuation and a higher trunk acceleration amplitude in OF, which implies a decreased cushioning (impact absorption) and hence less preservation of the head’s stability (Menz et al., 2003). Even though foot (vertical) accelerations were lower in OF, suggesting less impact, the OF were not able to attenuate the higher accelerations in the trunk. This reduction in impact absorption may be caused by age-related neuromuscular changes, such a reduced muscle strength of the triceps surae and quadriceps femoris(Reeves et al., 2006), degraded stiffness and elastic modulus of the tendons(McCrum et al., 2018), muscle co-contraction and degraded absorption of the intervertebral disc(Brzuszkiewicz-Kuźmicka et al., 2018). Considering that two-thirds of the weight of the human body is in the upper body, such higher trunk accelerations may be destabilizing, which may cause falls (Woollacott and Tang, 1997).

For stability, LDE calculated from trunk time-series data have been shown to better reflect differences in gait stability due to age than LDE calculated from data of other segments (Punt et al., 2015). In our study, OF showed significantly lower local dynamic stability (higher LDE) in ML acceleration, ML and VT angular velocity. Among these, the LDE calculated from trunk ML angular velocity had the largest effect size. As stability in the ML direction needs more control than stability in the AP direction during gait (Bruijn et al., 2009;O’Connor and Kuo, 2009), decreased LDE of trunk angular velocity in ML direction could be an early indicator of gait stability problems.

### 4.2 Variability measures

All participants in this study walked under the same environmental conditions. Thus, any between-subject differences in variability arose from differences in (internal) neuromotor noise and not (external) environmental noise. No differences were found in the variability of spatiotemporal measures, which was consistent with a previous study showing that temporal gait variability of older non-fallers was not significantly different from young adults in terms of standard deviation and coefficient of variation (Hausdorff et al., 1997).

Our OF walked with similar variability of maximum vertical acceleration of feet variability compared to YF (Table 3 and Supplementary Table 4). However, the variability of ML and VT acceleration amplitude of the trunk was larger for the OF, which could suggest OF are at a higher risk of balance loss and falling (Kavanagh and Menz, 2008).

All in all, our findings suggests that stability of the trunk might be a more sensitive indicator of locomotor impairment and potential future risk of falls than changes in variability of the trunk, as the LDE had higher effect sizes (Kang and Dingwell, 2008). Measures of variability of acceleration of the feet showed even higher effect sizes and might thus be even more useful. However, here, it should be noted that these effects were opposite from theoretically expected, with the OF having lower (means and variability) acceleration of the foot.

### 4.3 Limitations

All tests in our study were aimed at testing the same hypothesis, that is, OF are less stable and more variable than YF, hence, we did not use a correction for multiple testing. Nonetheless, not correcting may lead to Type I errors, and thus, some caution is warranted. Furthermore, the older participants in our study were quite fit and additional studies are needed to further investigate the applicability of acceleration attenuation when studying older adults. Future research can expand the sample size and conduct a multi-center study to obtain more representative results. Although we used only trunk and feet sensors for practical usefulness, the underlying mechanisms for the alterations in gait in the older women remain unclear and would require more detailed assessments of e.g. whole body kinematics and muscle activity.

## 5 Conclusions

Although healthy older females had similar walking speed and spatiotemporal parameters as young females during steady state walking, they showed lower vertical foot accelerations and higher vertical trunk accelerations, suggesting less impact, and less absorption of the impact. In addition, lower gait stability and higher variability of trunk movements for older females also indicated they were more likely to fall. The measures derived from the accelerations of the trunk were sensitive to reflect the gait instability as expected, especially trunk-foot vertical acceleration attenuation and its variability. While the variability of foot acceleration amplitudes was also sensitive to age, these differences were opposite from expected, making it harder to draw any conclusion as to their usefulness for fall prediction. These findings suggest that instrumented gait measurements may help for early detection of changes or impairments in gait performance, even before this can be observed by clinical eye or gait speed.

## Supporting information

see Supplementary Table 1, see Supplementary Tables 2 and 3 for dimension and delay values

## 6 Conflicts of Interest

The authors declare no conflict of interest.

## 7 Author Contributions

**Yuge Zhang:** Methodology, Software, Formal analysis, Investigation, Data curation, Writing-Original Draft, Writing-Review and Editing, Visualization. **Xinglong Zhou:** Conceptualization, Resources. **Mirjam Pijnappels:** Formal analysis, Writing review and Editing, Supervision, Funding acquisition. **Sjoerd M. Bruijn:** Methodology, Software, Formal analysis, Writing-Review and Editing, Supervision, Funding acquisition. All authors have read and agreed to the published version of the manuscript.

## 8 Acknowledgments

YZ was funded by a CSC Scholarship Council (CSC) fellowship (202009110145). MP funded by a VIDI Grant (no. 91714344) from the Dutch Organization for Scientific Research (NWO). SMB was funded by a VIDI grant (016.Vidi.178.014) from the Dutch Organization for Scientific Research (NWO).

## Notes

### Competing Interest Statement

The authors have declared no competing interest.

### Summary of Updates

This version corrected a small mistake.

