## Supplementary material for "Differences in gait stability and acceleration characteristics between healthy young and older females": see Supplementary Table 1, see Supplementary Tables 2 and 3 for dimension and delay values

**Table 1.** Coefficient of Variation (and SD) of all measures

| **Measures** | **OF** | **YF** | | **p-value** | **Effect Size** |
| --- | --- | --- | --- | --- | --- |
| **Spatial-temporal Gait Measures** |  |  |  | |  |
| Stride time | 0.02(0.005) | 0.02(0.004) | 0.86 | | -0.36 |
| IDS % | 5.28(0.90) | 5.11(0.84) | 0.56 | | -0.34 |
| TDS % | 5.67(1.70) | 5.58(0.86) | 0.83 | | -0.34 |
| Swing % | 2.10(0.42) | 2.11(0.57) | 0.94 | | -0.36 |
| Velocity of feet | 0.06(0.03) | 0.08(0.04) | 0.11 | | -0.64 |
| Stride length of feet | 0.02(0.01) | 0.03(0.03) | 0.16 | | -0.35 |
| **VT Acceleration Maximum (m/s^2^)** |  |  |  | |  |
| VT acceleration maximum of feet | 0.16(0.04) | 0.23(0.62) | 0.002* | | -1.40 |
| VT acceleration maximum of trunk | 0.06(0.02) | 0.05(0.01) | 0.13 | | 0.47 |
| **Trunk-foot vertical acceleration attenuation(m/s^2^)** | 0.24(0.05) | 0.27(0.05) | 0.94 | | -0.55 |
| **Acceleration Amplitude (m/s^2^)** |  |  |  | |  |
| AP acceleration amplitude of feet | 0.07(0.01) | 0.08(0.02) | 0.09 | | -1.38 |
| ML acceleration amplitude of feet | 0.15(0.03) | 0.21(0.07) | <0.001** | | -2.67 |
| VT acceleration amplitude of feet | 0.08(0.02) | 0.14(0.03) | <0.001** | | -0.76 |
| AP acceleration amplitude of trunk | 0.14(0.03) | 0.15(0.05) | 0.10 | | -0.12 |
| ML acceleration amplitude of trunk | 0.10(0.03) | 0.10(0.02) | 0.11 | | 0.22 |
| VT acceleration amplitude of trunk | 0.11(0.04) | 0.09(0.02) | 0.23 | | 0.50 |

Notes: AP = anteroposterior direction; ML = mediolateral direction; VT = vertical direction. P-values refer to group comparisons based on t-tests, except for measures of the feet, where they refer to the main effect of Group.

**Table 2.** Average (SD) of embedding dimension

| Time series | OF | YF |
| --- | --- | --- |
| AP acceleration of trunk | 12.10(5.88) | 11.88(4.97) |
| ML acceleration of trunk | 19.70(7.05) | 19.71(7.02) |
| VT acceleration of trunk | 24.75(7.57) | 26.29(7.97) |
| AP angular velocity of trunk | 11.10(3.46) | 10.47(3.78) |
| ML angular velocity of trunk | 14.35(6.98) | 15.00(3.48) |
| VT angular velocity of trunk | 9.45(3.17) | 12.06(2.61) |

**Table 3.** Average (SD) of delay time

| Time series | OF | YF |
| --- | --- | --- |
| AP acceleration of trunk | 5.95(0.94) | 4.76(1.30) |
| ML acceleration of trunk | 4.60(1.23) | 6.41(1.87) |
| VT acceleration of trunk | 4.20(2.17) | 3.76(1.75) |
| AP angular velocity of trunk | 7.05(1.82) | 8.24(2.41) |
| ML angular velocity of trunk | 5.90(1.48) | 6.18(1.42) |
| VT angular velocity of trunk | 9.90(3.54) | 8.53(1.91) |
